# Potent neutralizing equine antibodies raised against recombinant SARS-CoV-2 spike protein for COVID-19 passive immunization therapy

**DOI:** 10.1101/2020.08.17.254375

**Authors:** Luis Eduardo R. Cunha, Adilson A. Stolet, Marcelo A. Strauch, Victor A. R. Pereira, Carlos H. Dumard, Andre M. O. Gomes, Patrícia N. C. Souza, Juliana G. Fonseca, Francisco E. Pontes, Leonardo G. R. Meirelles, José W. M. Albuquerque, Carolina Q. Sacramento, Natalia Fintelman-Rodrigues, Tulio M. Lima, Renata G. F. Alvim, Federico F. Marsili, Marcella Moreira Caldeira, Luisa M. Higa, Fábio L. Monteiro, Russolina B. Zingali, Guilherme A. P. de Oliveira, Thiago M. L. Souza, Amilcar Tanuri, Andréa C. Oliveira, Herbert L. M. Guedes, Leda R. Castilho, Jerson L. Silva

## Abstract

We used the trimeric spike (S) glycoprotein (residues 1-1208) in the prefusion conformation to immunize horses for production of hyperimmune globulins against SARS-CoV-2. Serum antibody titers measured by anti-spike ELISA were above 1:1,000,000, and neutralizing antibody titer was 1:14,604 (average PRNT_90_), which is 140-fold higher than the average neutralizing titer of plasma from three convalescent COVID-19 patients analyzed for comparison. Using the same technology routinely used for industrial production of other horse hyperimmune products, plasma from immunized animals was pepsin digested to remove the Fc portion and purified, yielding a F(ab’)_2_ preparation with PRNT_90_ titers 150-fold higher than the neutralizing titers in human convalescent plasma. Repeating the hyperimmunization in a second group of horses confirmed the very high neutralizing titers in serum and in a GMP clinical F(ab’)_2_ lot. Virus-neutralizing activity in samples from mice that received the F(ab’)_2_ preparation was detected even three days after injection, indicating an appropriate half-life for therapeutic intervention. These results supported the design of a clinical trial (identifier NCT04573855) to evaluate safety and efficacy of this horse F(ab’)_2_ preparation.

## Introduction

The pandemic caused by SARS-CoV-2, the etiological agent of COVID-19, is an urgent health problem worldwide, especially in the Americas (https://covid19.who.int/). Consequences for human health and for the global economy have been devastating. Considering the absence of approved antiviral treatments and vaccines and the uncertainties regarding antibody responses in individuals infected by SARS-CoV-2^1–4^, the COVID-19 scenario seems to be still far from a solution. Passive immunization using plasma from COVID-19 convalescent patients has been used as an alternative therapy^5,6^. However, the heterogeneous neutralizing titers in different convalescent donors and the simultaneous transfer of other plasma components are drawbacks that hinder its wide use.

The development of virus-neutralizing purified hyperimmune globulins produced in horses or llamas may be an approach to treat SARS-CoV-2 infection. The use of llamas^7^ to develop passive immunization therapies is still experimental and limited by animal availability. On the other hand, hyperimmune serum, immunoglobulins or IgG fragments produced in horses have been used to treat many diseases, such as rabies, tetanus and snake envenomation, among others^8,9^.

Brazil, such as many other countries, has a large established capacity to produce equine hyperimmune globulin preparations for a range of indications, which makes the production of such products against SARS-CoV-2 highly feasible. Previous works with other related betacoronaviruses reported that equine hyperimmune sera resulted in neutralizing antibodies against SARS-CoV^10^ and MERS-CoV^11^. In these works, immunization was performed using virus or virus-like particles, using complete Freund’s adjuvant (CFA) in the first immunization and incomplete Freund’s adjuvant (IFA) in the subsequent immunizations. More recently, the recombinant receptor-binding domain (RBD) of the SARS-CoV-2 S protein was shown to stimulate antibody production in mice and equines^12,13^.

In the present work, we also used a recombinant antigen to immunize horses. However, the antigen chosen was the trimeric version of the complete ectodomain of the spike protein, comprising both the S1 (responsible for receptor binding) and the S2 (responsible for fusion to the cell membrane) domains. Moreover, the chosen antigen was produced in house using a gene construct that yields the protein in a stabilized, trimeric prefusion conformation^14^, with the aim of maximizing the formation of high-quality neutralizing antibodies. Our strategy may be easily reproduced in any part of the world and could be rapidly tested as a therapy for COVID-19. Equine immunization is a well-known and easily scalable technology proven for generating high titers of neutralizing antibodies, thus showing advantages over other strategies, such as using convalescent human plasma. In this study, we demonstrated extremely high neutralizing titers obtained by means of equine immunization. The final purified F(ab’)_2_ preparation had an average PRNT_90_ neutralizing titer 150-fold higher than that of human convalescent plasma from three patients in Brazil. A GMP clinical lot has been produced for use in a Phase 1 clinical trial (identifier NCT04573855).

## Results

### Immunization with trimeric S protein induces high titers of specific equine IgG

We produced trimers of the spike protein in the prefusion conformation^14^ by stably expressing the gene construct in mammalian cells as described previously^15^. A quality check of the spike protein was performed by SDS-PAGE, and its trimeric prefusion conformation was verified by size-exclusion chromatography (Fig. S1). Initially, five horses were immunized with six subcutaneous injections of S protein, with an interval of one week between inoculations. No adverse effects or animal suffering were observed for any of the five horses that received the protein injections. Anti-spike IgG measured by enzyme-linked immunosorbent assay (ELISA) in weekly serum samples showed that one week after the first immunization anti-SARS-CoV-2 IgG was not yet detectable but that IgG titers increased progressively after successive immunizations (Fig. 1A). Four out of the five immunized horses produced similar amounts of specific antibodies, but one did not show a strong response (Fig. 2). Despite the low-responding horse (#835), the average IgG titer for all five horses reached 1,180,980 after 42 days, i.e., one week after the sixth immunization (Fig. 1B), indicating that the trimeric S protein is a good immunogen to induce the production of specific anti-SARS-CoV-2 antibodies.

**FIGURE 1.**
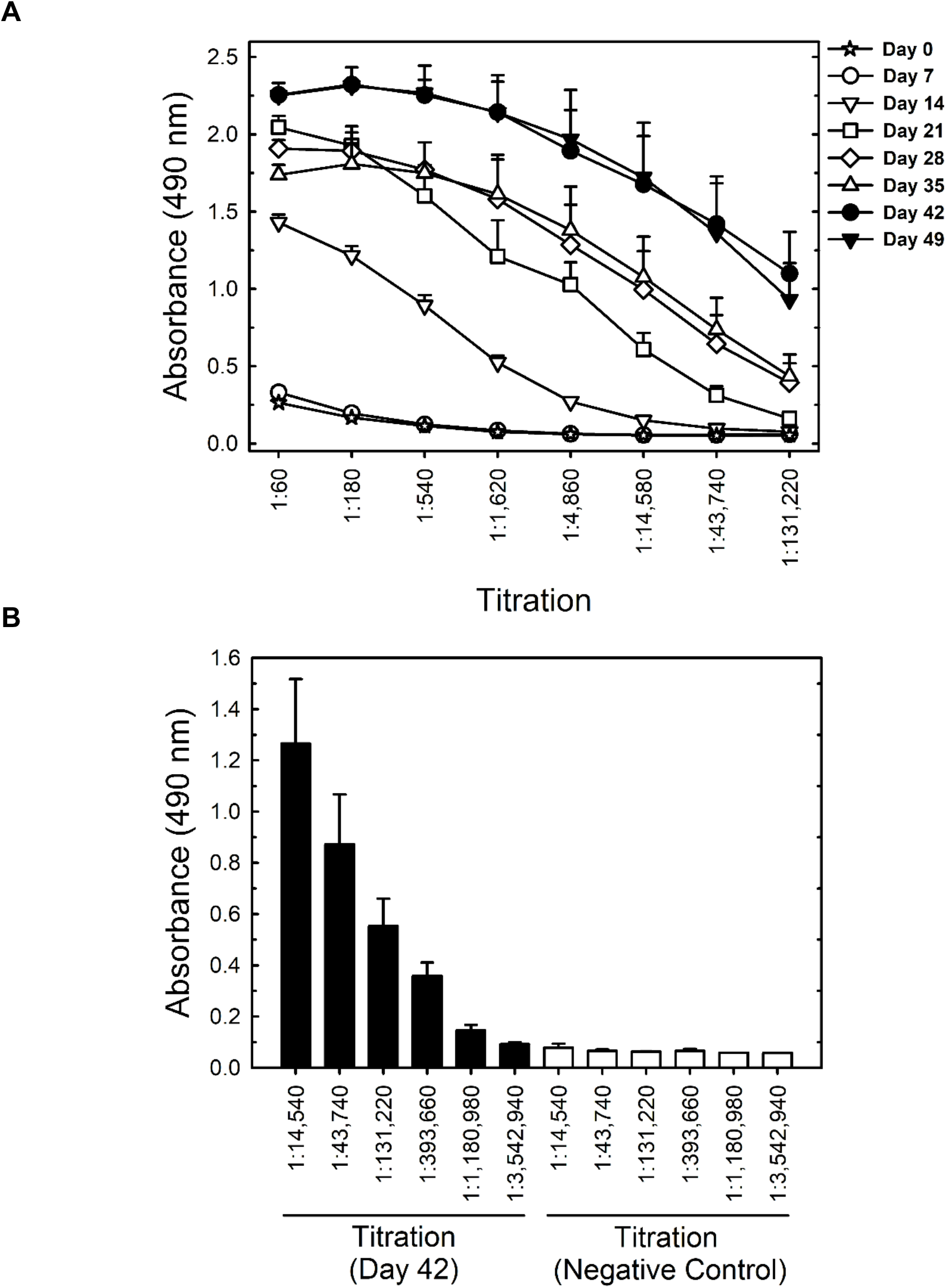
Titration of horse antibodies by S protein ELISA in samples from the first group of animals. **(A)** Titration up to 131,200-fold dilution of serum samples collected from immunized horses on different days after initial immunization. **(B)** Titration up to a 3,542,940-fold dilution of samples collected on day 42 after the first immunization. All results shown are the mean ± standard error for all 5 horses. The negative control is a pool of preimmune sera collected from all 5 horses of Group 1.

**FIGURE 2.**
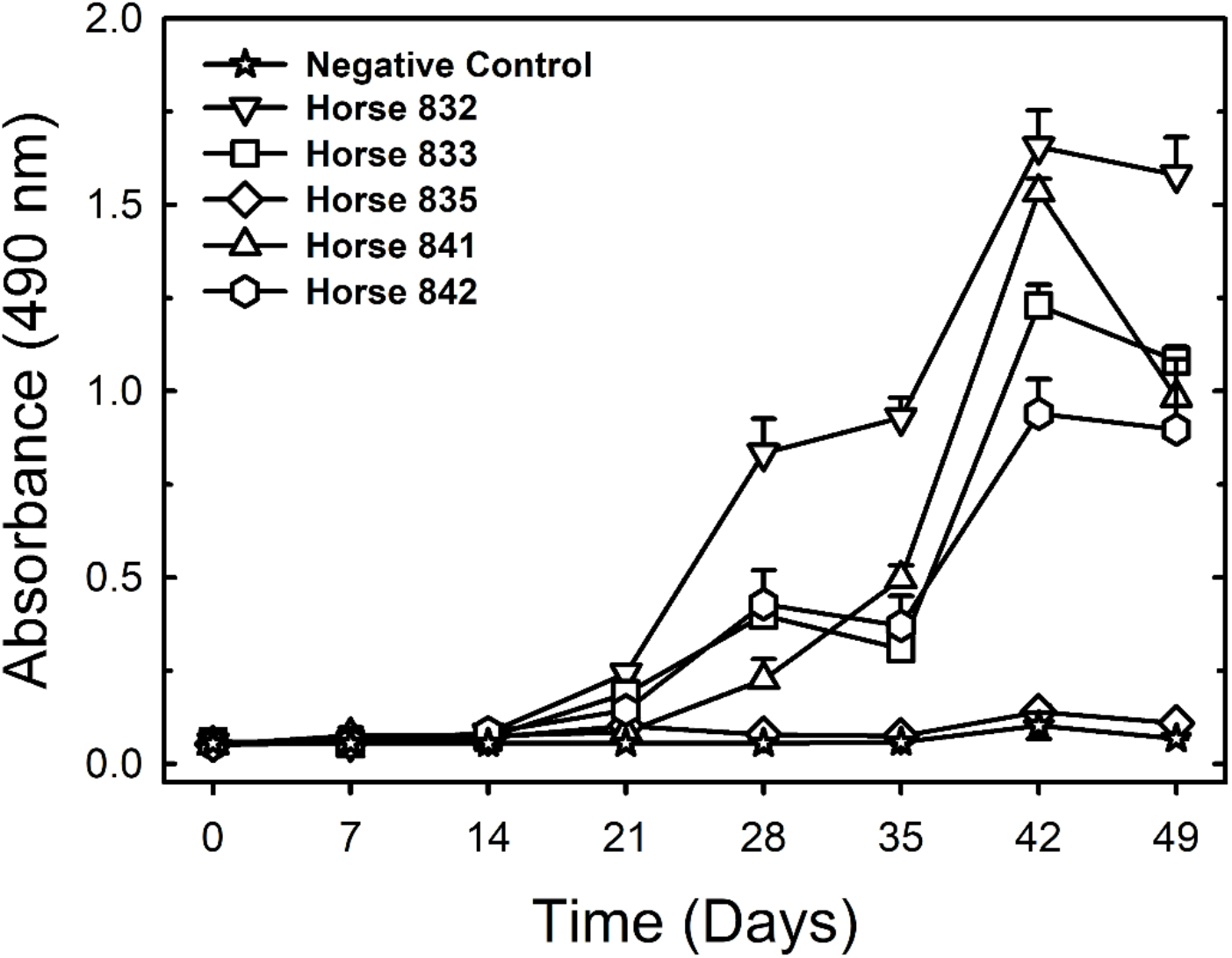
Anti-S antibodies measured by ELISA in serum samples collected over time (first group of animals). All sera were diluted 1:131,220. The results are shown as the mean ± standard error for analytical duplicates. The negative control is a pool of preimmune sera collected from all 5 horses of Group 1.

### Equine sera and F(ab’)_2_ fragments developed against trimeric S protein have potent neutralizing titers against SARS-CoV-2

We first evaluated the *in vitro* neutralizing activity against SARS-CoV-2 of equine sera collected after four to six immunizations (days 28 to 49 after the first immunization). PRNT_50_ titers (average values for all five horses) seemed to achieve a plateau of approximately 1:23,000 after the fourth immunization, whereas the more stringent PRTN_90_ titers (average values for all five horses) still showed an ascendant trend over time, reaching an average PRNT_90_ titer of 1:14,604 on day 49. Due to the low-responding horse (#835), the standard deviations were high (Fig. 3; Table S1).

**FIGURE 3.**
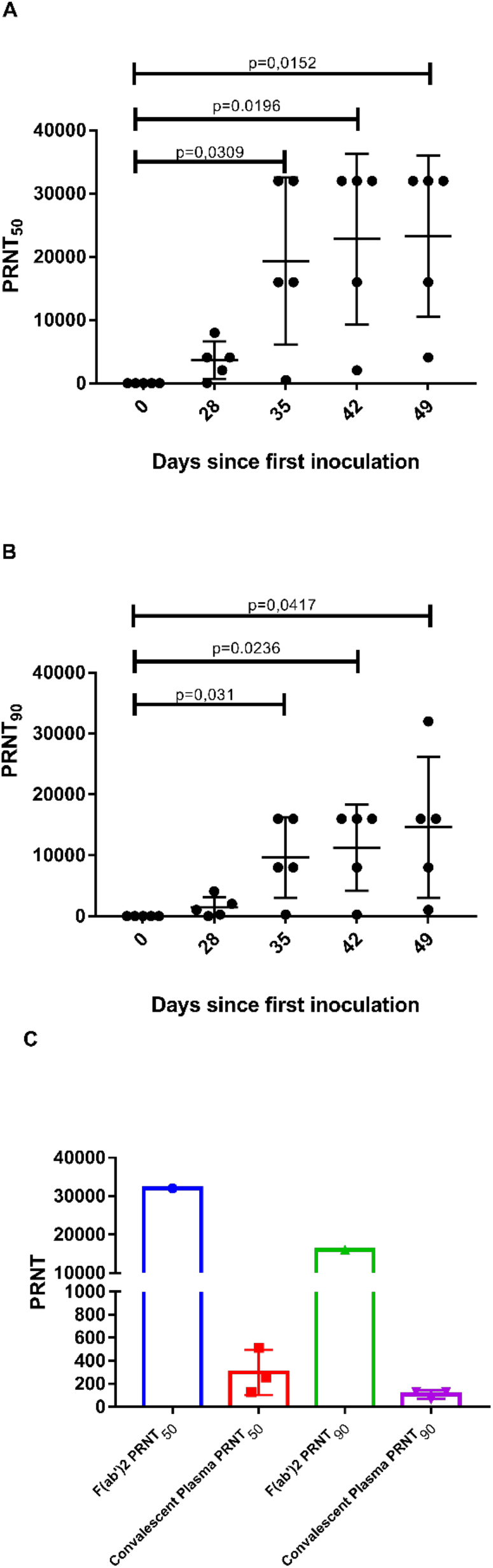
Microneutralization assays. **(A)** PRNT_50_ and **(B)** PRNT_90_ for equine plasma collected on days 28 to 49 after the first immunization. Data shown as the mean ± SD; p<0.05 vs plasma collected on day 0 according to a t-test for paired samples. **(C)** Comparison of PRNT_50_ and PRNT_90_ for equine F(ab’)_2_ concentrate and human convalescent plasma. Data is shown as mean ± standard deviation.

Plasma from all five horses was then pooled, digested with pepsin to cleave off the Fc portion and precipitated with ammonium sulfate for purification, resulting in a concentrate of F(ab’)_2_ fragments containing approximately 90 mg/mL total protein. The F(ab’)_2_ concentrate maintained the capacity to recognize the trimeric S protein, displaying an ELISA titer of 1:1,000,000 (Fig. 4). In order to confirm if F(ab’)_2_ fragments also maintained their capacity to recognize SARS-CoV-2 proteins in cell culture, we prepared a F(ab’)_2_-FITC conjugate, which was shown to specifically bind to SARS-CoV-2 infected Vero E6 cells (Figure S2). F(ab’)_2_ neutralizing titers were also very high, achieving a PRNT_50_ of 1:32,000 and a PRNT_90_ of 1:16,000 (Fig. 3 and Table S1).

**FIGURE 4.**
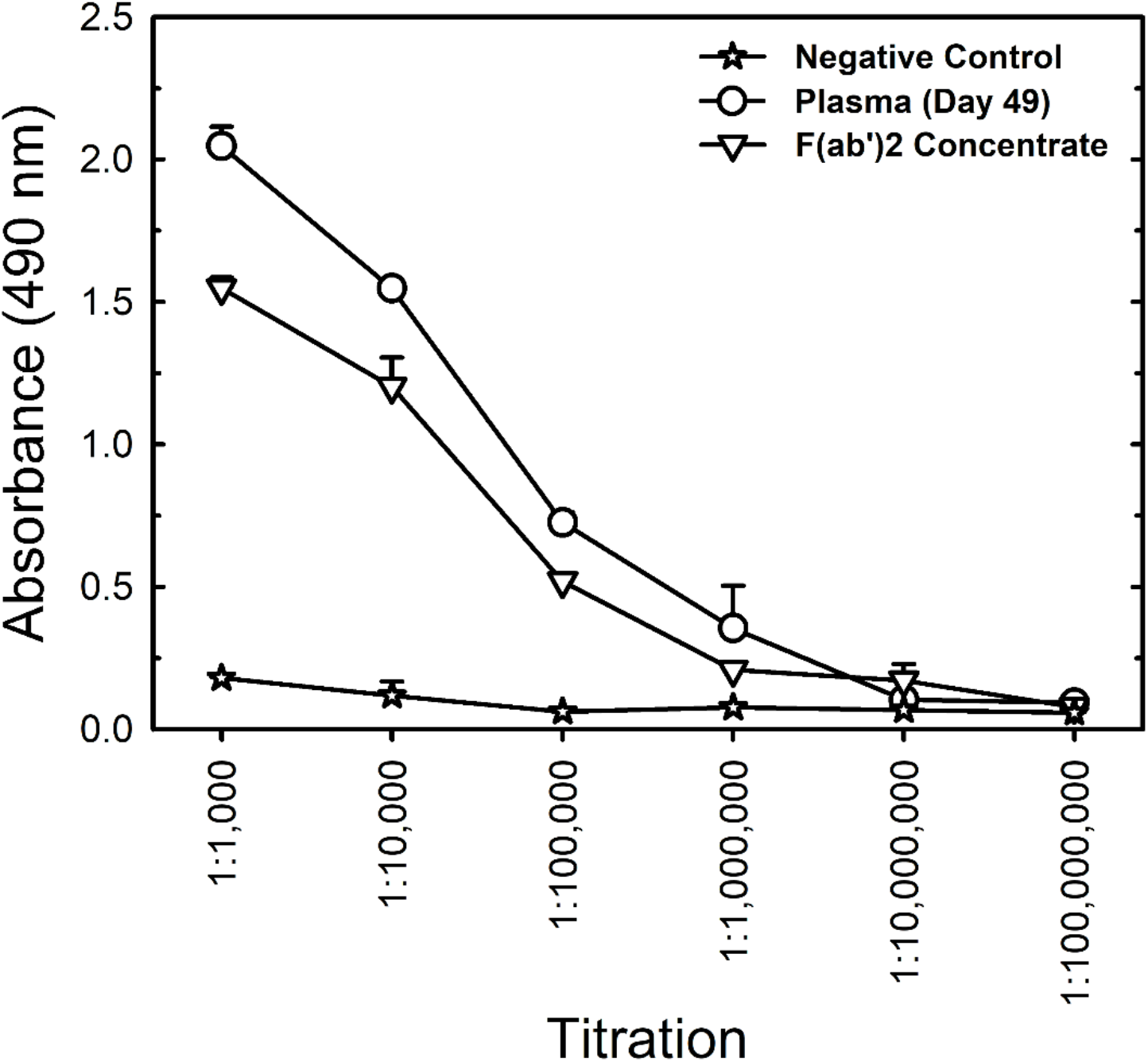
Comparison of S protein ELISA titration of equine F(ab’)_2_ concentrate and horse plasma collected on day 49 after the first immunization of the first group of animals. The sample from day 49 and the preimmune negative control are pools of samples collected from all 5 horses of Group 1. The results are shown as the mean ± standard error for analytical duplicates.

We further compared the neutralizing titers of the equine samples to the neutralizing titers determined for plasma from three convalescent COVID-19 patients. Interestingly, the average neutralizing titers of equine serum (from day 49) were 78- and 138-fold higher than the average human convalescent plasma titers in terms of PRNT_50_ and PRNT_90_, respectively. Regarding the F(ab’)_2_ concentrate, the neutralizing titers were 107-fold (for PRNT_50_) and 151-fold (for PRNT_90_) higher than the average human convalescent plasma titer. These data show the great potential of using equine immunoglobulins in the treatment of COVID-19.

### Sustainable production of potent antibodies against SARS-CoV-2

In order to evaluate sustainability of equine antibody production and to enable scale-up to industrial manufacturing, the first group of five horses was reimmunized, and a second group of five horses was immunized using the same protocol as the first group (Figure 5). Reimmunization of the first group occurred by means of three spike protein injections on weeks 13, 14 and 15, i.e. 6 weeks after plasma collection on day 49 (week 7). The titer of antibodies, which had decreased since the first bleeding, was boosted and reached levels slightly higher than after the first series of immunizations (Fig. 5A). The time interval to boost and the decreased number of immunizations to boost is a standard protocol applied to the production of other equine hyperimmune sera produced at IVB.

**FIGURE 5.**
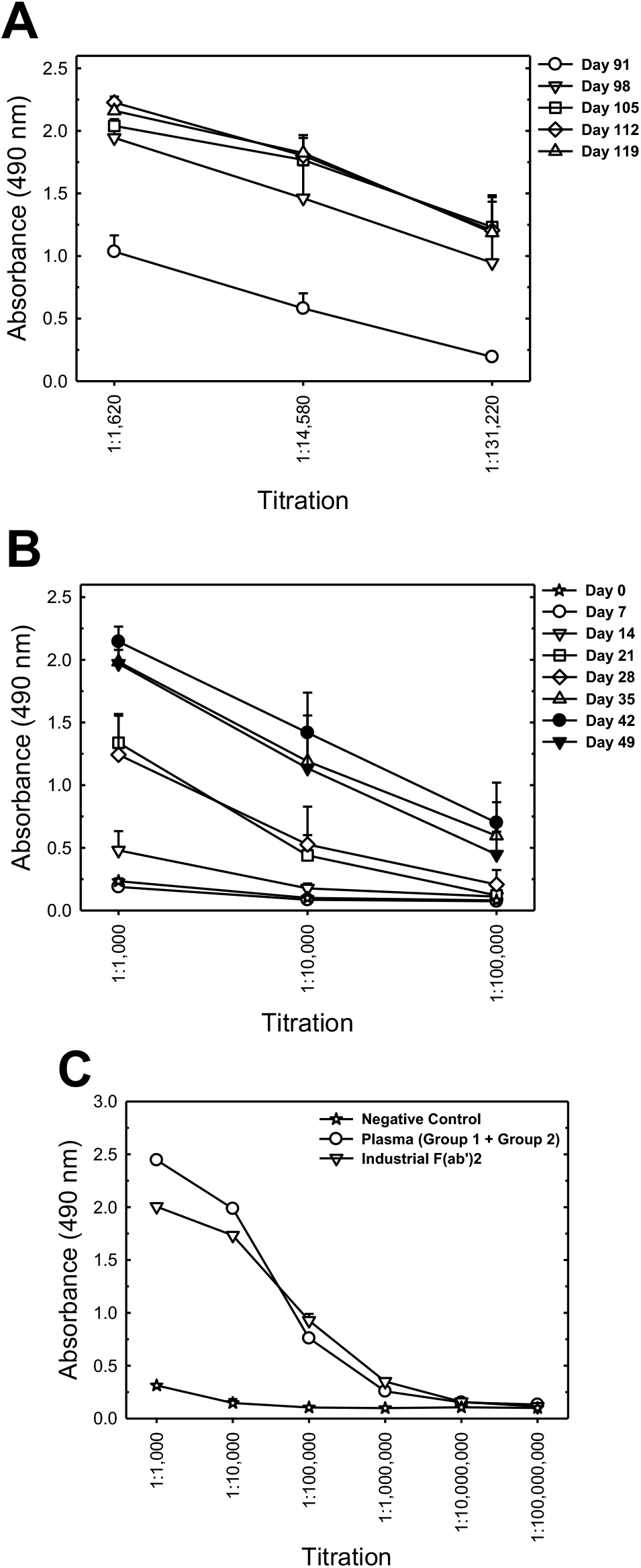
S protein ELISA titration of Group 1 reimmunization, Group 2 (first immunization) and industrial F(ab’)_2_ concentrate. **(A)** Titration of pooled sera from all 5 horses of Group 1 upon reimmunization (injections on days 91, 98 and 105). **(B)** Titration of pooled sera from all 5 horses of Group 2 (first immunization: 6 weekly injections on days 0-35). **(C)** Titration of the pool of plasma collected from Groups 1 and 2 on day 49 and the industrial F(ab’)_2_ concentrate manufactured under GMP conditions from this pool of plasma. Data is shown as mean ± standard error of analytical duplicates. The negative control is a pool of preimmune sera of Group 1 animals.

The immunization scheme applied to the second group of five horses was the same as for the first immunization of the first group of animals (6 weekly spike protein injections) and equally resulted in increasing anti-S antibody titers (Fig. 5B).

The plasma collected in the first bleeding of the first group of horses and in the first bleeding of the second group of horses was pooled and resulted in 160 L of plasma that was processed in the industrial facility of Vital Brazil Institute (IVB), which is certified for good manufacturing practices (GMP), resulting in the first GMP-compliant lot of anti-COVID-19 equine F(ab’)_2_ active pharmaceutical ingredient (API). Fig. 5C shows a comparison of ELISA titers of this GMP F(ab’)_2_ concentrate lot and of the plasma pool used as raw material for its production. Antibody titers were in the range of 1:1,000,000 as obtained for the pilot lot produced from the first bleeding of the first horse group (Fig. 4).

Neutralizing activity in the GMP API (Table S1) was 1:65,556 as PRNT_50_ and 1:16,384 as PRNT_90_, showing that the well-established routine GMP process implemented at IVB leads to higher F(ab’)_2_ yields than the pilot process using laboratory equipment. This API was further processed (dilution in water-for-injection, fill and finish) for use in the Phase I/II clinical trials of the anti-COVID-19 F(ab’)_2_ concentrate.

### Persistence of neutralizing activity after F(ab’)_2_ injection in mice

To test the clearance of the neutralizing activity, mice were injected with different doses (10, 25 or 50 μg) of the pilot lot of equine F(ab’)_2_ concentrate. After 72 h, blood was collected, and neutralizing activity in mice plasma was determined (Fig. 6). The data clearly shows that high neutralizing titers are maintained for at least 3 days after injection, with PRNT_50_ and PRNT_90_ titers for the highest dose (50 μg) being comparable to human convalescent plasma (Table S1). This result indicates that the F(ab’)_2_ fragments would have a sufficient half-life to reach different organs, including the lungs, to exert its antiviral activity.

**FIGURE 6.**
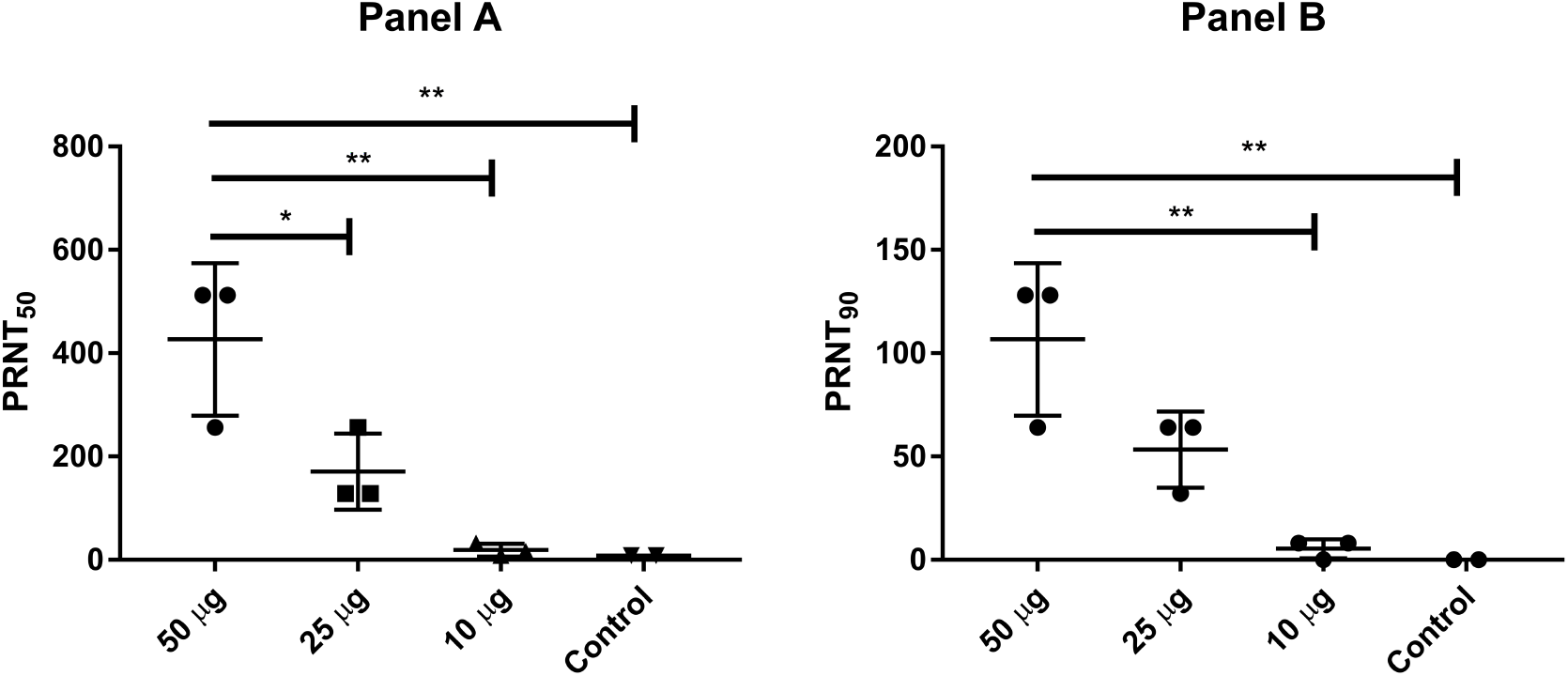
Mice injected with equine F(ab’)_2_ still presented a neutralizing response in plasma 3 days post injection. Mice were injected via intraperitoneal inoculation with equine F(ab’)_2_ (50 μg, 25 μg or 10 μg total protein concentration), and blood was collected 72 h after injection. Neutralizing titers were determined by PRNT assay. The PRNT_50_ and PRNT_90_ results are shown in panels A and B, respectively. Data are shown as the mean ± SD; *=p<0.05; **=p<0.01 according to one-way ANOVA and Tukey’s post hoc test.

## Discussion

In the setting of a pandemic, when no vaccines and no specific treatments are available, passive immunotherapies using convalescent human plasma or animal-derived hyperimmune globulins usually represent the first specific antiviral therapies to become available. In previous outbreaks caused by other viruses, such as SARS-CoV, MERS-CoV, Ebola and avian influenza virus, horse immunization to produce hyperimmune globulins was evaluated^10,11,16,17^. In addition to their application in emerging and reemerging infectious diseases, heterologous polyclonal antibody therapies are also useful to treat longstanding neglected tropical diseases^18^. Equine antivenom products are routinely produced in both high- and low-income countries^16^, and WHO standardized guidelines are available for this class of products (www.who.int/bloodproducts/snake_antivenoms/snakeantivenomguide/en), potentially allowing equine antibody products to be largely available worldwide within a relatively short period of time.

Motivated by the SARS-CoV outbreak in Asia in 2002-2003, equine F(ab’)_2_ was developed by immunizing horses 4 times using whole virus (the F69 strain propagated in Vero cells) and CFA/IFA, resulting in ELISA titers of 1:14,210 and neutralization titers of 1:14,240^10^. In the case of MERS-CoV, equine immunoglobulin products were generated by immunizing horses with virus-like particles (VLPs) formed by three viral structural proteins (the spike, membrane and nucleocapsid proteins)^11^. The equine antibodies were then purified by immunoaffinity chromatography using a resin containing the RBD of the S protein as an affinity ligand^11^. The purified anti-MERS-CoV antibody preparation presented ELISA titers of 1:20,480 and neutralization titers of 1:20,900. The binding and neutralizing titers obtained in the present work are quite superior (1:1,000,000 and 1:23,219, respectively) and indicate a high immunogenicity of the trimeric S protein used herein.

Recently, two reports have described the production of equine antibodies by immunizing horses with the recombinant SARS-CoV-2 RBD^12,13^, which comprises approximately 16% of the spike protomer. Pan et al. evaluated a 4-dose immunization scheme in 4 horses, totaling 33 mg of RBD injected per horse over 28 days since the first immunization^12^, whereas Zylberman et al. evaluated a total of 3.5 mg of RBD per horse in 2 horses, also for 4 immunizations over 28 days^13^. Both works reached titers of binding antibodies in the range of 1:1,000,000, similar to the titers obtained in the present work. Zylberman et al. apparently achieved complete neutralization at a 1:10,240 dilution^13^, whereas Pan et al. reported 80% neutralization for a 1:2,560 dilution^12^. In the present work, although our immunization strategy used only 1.2 mg of S protein per each of the 5 horses (6 weekly doses of only 200 μg), we achieved binding titers over 1:1,000,000 (Fig. 1B), and more importantly, average titers for 90% neutralization were 1:14,604 when including the low-responding horse or 1:18,000 if the low-responding animal was not included (Table S1). The higher ratio of neutralizing to binding antibodies obtained in the present work indicates a high quality of the antibodies generated by using the prefusion trimeric spike protein as the immunogen. Our results are in agreement with a previous comparison of mice immunized with the trimeric spike protein and the RBD, which showed higher neutralizing titers for the S protein^19^. This finding is probably related to the fact that other domains of the spike protein besides the RBD are also targets for neutralizing antibodies^20^. Not surprisingly, most COVID-19 vaccines under development worldwide focus on the whole spike protein, either as the recombinant protein itself or as nucleic acid encoding the complete S protein, such as in the case of mRNA or vectored vaccines.

In our study, we used Montanide ISA 50V as an adjuvant. In previous studies, horse hyperimmunization against SARS-CoV^10^, MERS-CoV^11^ and SARS-CoV-2^12,13^ was performed using CFA/IFA. The Montanide adjuvant enhances the immune response by warranting a depot effect and promoting slow delivery of the protein to maintain recognition, stimulation and phagocytosis by dendritic cells and B cells for long time periods^21^. The present results not only indicate the high immunogenicity of the trimeric prefusion spike protein but also demonstrate the feasibility of using Montanide, which is a less reactogenic adjuvant than CFA/IFA^22^.

To avoid the risk of antibody-dependent enhancement (ADE) of infection, the pooled plasma obtained from the horses was processed to remove the Fc portion of IgG by pepsin digestion, followed by partial purification of the F(ab’)_2_ fragment by ammonium sulfate fractionation. This method is a proven technology used worldwide for the production of equine hyperimmune products, since the use of F(ab’)_2_ instead of whole IgG eliminates nonspecific binding between the Fc portion of antibodies and Fc receptors on cells, thus avoiding ADE. This technology is regularly used for GMP manufacturing of anti-rabies, anti-tetanus and anti-venom F(ab’)_2_ products at Vital Brazil Institute (IVB), with no records of hypersensitivity issues and decades of excellent safety records. Additionally, safety trials of an analogous equine F(ab’)_2_ product against avian influenza (H5N1) performed by Bal et al. confirmed that it is a well-tolerated product, causing just a few mild adverse events^17^. Importantly, the anti-SARS-CoV-2 F(ab’)_2_ concentrate developed herein maintained the very high binding and neutralizing titers found in unprocessed sera (Figs. 3, 4 and 5, and Table S1).

As mentioned above, convalescent plasma is another therapeutic alternative that quickly becomes available in the setting of outbreaks. For comparison, we evaluated the neutralization titer of three human convalescent plasma samples. We found that the neutralization titers of the equine F(ab’)_2_ concentrates were two orders of magnitude higher than those of human convalescent plasma (107- and 151-fold higher for PRNT_50_ and PRNT_90_, respectively). Plasma from the low-responding horse, #835, which started to show relevant neutralizing activities just after the fourth inoculation, was included in the plasma pool used to produce the pilot-scale F(ab’)_2_ concentrate. Thus, if horses were preselected for their neutralizing activity prior to F(ab’)_2_ manufacture, the neutralizing ability of the antibody concentrate could be even higher, reducing infusion volumes in patients. Zylberman et al., who used SARS-CoV-2 RBD to immunize horses, reported that their F(ab’)_2_ product had an approximately 50-fold higher neutralizing capacity than convalescent plasma reported in the literature^13^. In the other work using the RBD to immunize horses evaluated samples from 11 COVID-19 convalescent patients and observed that at a 1:640 dilution, three human samples showed 50% neutralization, and another two showed 80-90% neutralization ability. In contrast, their unprocessed horse antisera after three immunizations showed complete neutralization at a 1:640 dilution and 50-60% neutralization at a 1:10,240 dilution^12^. Overall, these data confirm the promising potential of equine hyperimmune products as COVID-19 countermeasures over human convalescent plasma, considering the high potency and safety record of F(ab’)_2_ products as well as eventual limitations related to human plasma availability and eventual risks of adventitious agent transmission by human plasma products.

The advantage of using the fully folded trimeric S protein in the prefusion conformation to elicit equine polyclonal antibodies against SARS-CoV-2 is the main novelty of our work. In a recent report, Liu et al. reported that highly neutralizing monoclonal antibodies discovered from B cells of convalescent COVID-19 patients targeted epitopes in the RBD and in the N-terminal domain (NTD), as well as quaternary epitopes of the trimeric spike protein^20^. Thus, the use of the trimeric spike protein combines the advantages of higher immunogenicity than smaller protein fragments (such as the RBD used in other works^12,13^) with low biosafety concerns compared to those that would apply in case of inoculating the whole virus (as used for SARS-CoV^10^). Fig. 7 illustrates our whole strategy and emphasizes the potential binding of F(ab’)_2_ fragments to the S protein on the viral surface. The presence of neutralizing activity in mice three days after F(ab’)_2_ injection (Fig. 6) supports its use for passive immunization treatment of hospitalized patients. Because multiple organs can be affected by SARS-CoV-2, resulting in a worse prognosis, the relatively long half-life of F(ab’)_2_ is likely to prevent disease progression.

**FIGURE 7.**
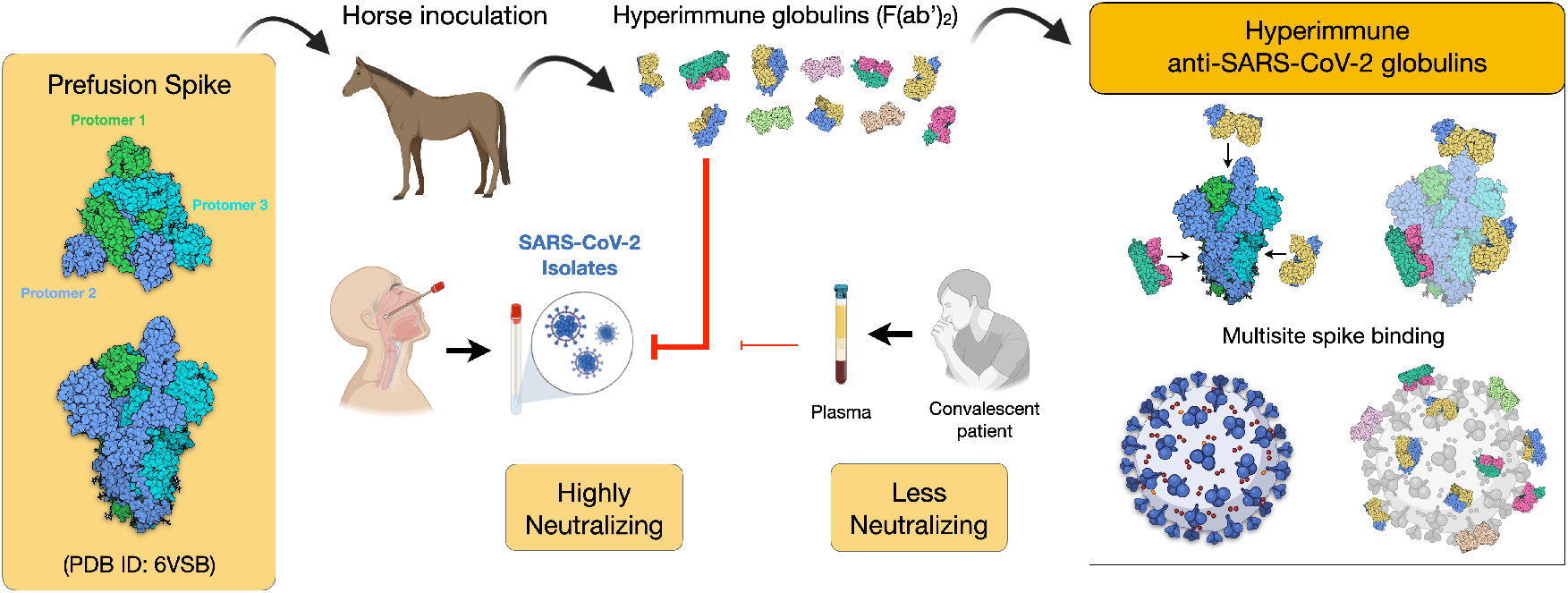
Scheme of the immunization strategy and anti-SARS-CoV-2 hyperimmune globulin production. Recombinant prefusion trimeric S protein is used to inoculate horses and to produce hyperimmune F(ab’)_2_ concentrate. The equine antibody preparation presented a much higher capacity to neutralize a SARS-CoV-2 isolate than human convalescent plasma. One advantage of using the full-length recombinant spike trimer is the production of antibodies against different antigenic segments of the viral protein. This strategy may result in more efficient neutralizing capacity than that of antibodies produced against isolated fragments of the spike protein, such as the RBD.

In summary, the use of an anti-COVID-19 hyperimmune F(ab’)_2_ concentrate produced by using a potent antigen (the prefusion trimeric spike glycoprotein) is a prompt alternative for passive immunization therapy, especially in a situation in which no vaccine has been approved (Fig. 7). Even in a more optimistic scenario of efficacious vaccine(s) becoming available, the possibility of having equine antibody products is highly important, as has been proven in the case of the rabies vaccine, where both passive and active immunizations are used to save lives.

Hyperimmune F(ab’)_2_ concentrates are manufactured in existing facilities available worldwide, in both high- and low-income countries, using a long-proven platform technology, which could help accelerate the regulatory pathway to product approval for human use. We have started the regulatory submission to perform a phase I/II clinical trial for safety and efficacy evaluation of treatment with anti-SARS-CoV-2 equine immunoglobulin (F(ab’)_2_) in hospitalized patients with COVID-19 in early stages of the disease, i.e. not requiring invasive ventilation support (ClinicalTrials.gov identifier: NCT04573855).

Taking as reference the production of other registered (Fab’)_2_ concentrates regularly produced by IVB in Brazil (e.g., anti-rabies immunoglobulin), each horse can supply at least 10 L of plasma for processing, resulting in at least 200 ampoules of F(ab’)_2_ concentrate. Since one horse can be bled up to six times per year without any animal suffering, this would mean that producing 100,000 ampoules (doses) per year would require approximately 80 horses per year. This is still a rough order-of-magnitude estimate, since the precise dose size still needs further studies to be defined.

Altogether, our results show that equine anti-COVID-19 immunoglobulin has a high chance of treating patients and contributing to changing the course of the COVID-19 pandemic.

## Methodology

### Production and purification of recombinant SARS-CoV-2 spike (S) glycoprotein

The cell line HEK293-COV2-S was generated by Alvim et al. (2020)^15^ and stably expresses the soluble ectodomain of the spike protein of SARS-CoV-2 in the prefusion trimeric conformation^14^. The cells were cultivated in HEK-GM medium (Xell AG, Germany), either at 300-mL scale at 37°C and 5% CO_2_ in Erlenmeyer flasks under orbital agitation (180 rpm, 5-cm stroke) or in 1.5-L stirred-tank bioreactors operated at a pH setpoint of 7.1 and a dissolved oxygen setpoint of 40% air saturation. Cell-free supernatant was obtained by microfiltration with 0.45-μm PVDF membranes and injected into a 5-mL StrepTrap XT affinity chromatography column (Cytiva) following the manufacturer’s instructions. Protein concentration, purity and identity in the eluted fractions were confirmed by NanoDrop (ThermoFisher), SDS-PAGE and Western blot analyses, respectively. The purified protein obtained in the affinity chromatography eluate was used to immunize horses and as antigen in the ELISA to detect anti-SARS-CoV-2 antibodies in samples.

### Animal immunization

All the procedures involving animals were performed in accordance with the animal research ethical principles determined by the National Brazilian Law 11.794/08. The protocol was approved by the Animal Care and Use Committee from IVB under permission no. 003.

All animals were subjected to prophylactic vaccination and deworming programs routinely utilized at the IVB farm and were tested for equine infectious anemia and *Burkholderia mallei*, as determined by the Brazilian Ministry of Agriculture regulations.

Five 3- to 5-year-old healthy horses (3 males and 2 females) from the IVB farm, weighing approximately 350 kg each, were used for the production of polyvalent sera. Before immunization, blood samples were drawn from the jugular vein, and sera were stored at −20°C for use as negative controls in the binding and neutralizing antibody determinations. Each horse was subcutaneously immunized six times at 7-day intervals (on days 0, 7, 14, 21, 28 and 35) in different positions of the dorsal region. Each immunization of each horse consisted of 200 μg of recombinant SARS-CoV-2 trimeric S protein mixed with Montanide ISA 50V adjuvant (Seppic, France) to form an emulsion (one part immunogen in sterile saline to one part sterile adjuvant). The horses were checked daily, and food and water intakes were monitored. Blood samples were collected every 7 days just before the next inoculation and 7 and 14 days after the last inoculation (up to day 49 since the first inoculation). Sera were stored at −20°C until the measurement of binding and neutralizing antibody titers.

### Production of bench-scale and industrial (GMP) F(ab’)_2_ lots

For the production of the bench-scale pilot lot, plasma processing to produce F(ab’)_2_ was initiated by adding 3 L of water and 15 mL of 90% phenol solution to 2 L of horse plasma in a reactor. The solution was homogenized for 10 min, and the pH was adjusted to 4.3. Under agitation, 1.25 g/L pepsin was added, the pH was adjusted to 3.2, and the sample remained under agitation for 10 min. The sample was then stirred at 37°C, and the pH was adjusted to 4.2 with sodium hydroxide. Under constant agitation, sodium pyrophosphate decahydrate (12.6 mM) and toluene (10 μM) were added. Later, ammonium sulfate was added at 12% (m/v), and the solution was incubated at 55°C for 1 h.

To separate the F(ab’)_2_ fragments, the Fc portion was precipitated and subsequently filtered under pressure at constant agitation. F(ab’)_2_ was recovered from the liquid phase. To subsequently precipitate F(ab’)_2_ from the liquid phase, ammonium sulfate was added at 19% (m/v), and a second precipitation was performed under constant agitation and alkaline pH. Subsequently, the solution was diafiltered using a 30-kDa tangential ultrafiltration system until ammonium sulfate became undetectable in the retentate. The samples were isotonized with 15 mM NaCl, and 90% phenol solution was added to a final concentration of 0.3% (v/v). After sterile filtration, the F(ab’)_2_ product was stored at 4°C.

The industrial scale GMP-compliant lot was produced according to an analogous process (but scaled up to start from 160 L of plasma), using the industrial GMP facility of IVB in Rio de Janeiro state, Brazil.

### Enzyme-linked immunosorbent assay (ELISA)

In brief, polystyrene high-adsorption 96-well microplates (ThermoFisher, USA) were coated with 500 ng/well recombinant SARS-CoV-2 S protein (100 μL/well at 5 μg/mL) in carbonate-bicarbonate buffer (pH 9.6) overnight at room temperature and then blocked with 3% BSA (Sigma, USA) in PBS for 2 h at 37°C. Serially diluted serum samples of the 5 horses were added to the plate and incubated at 37°C for 1 h. Horseradish peroxidase-conjugated rabbit anti-horse IgG (Sigma A6917, USA) diluted 1:10,000 in PBS was incubated at 37 °C for 1 h. After each step, the plates were washed 3 times with PBS containing 0.05% Tween 20 (PBST). OPD substrate (100 μL/well - Sigma, USA) was added to wells and incubated in the dark for 10 min at room temperature. The reaction was stopped by adding 50 μL of 30% H_2_SO_4_ (v/v) to each well, and the absorbance value was measured at 490 nm in a microplate reader (Epoch/2 microplate spectrophotometer, Biotek). The IgG antibody titer was defined as the highest dilution of serum yielding an absorbance ratio greater than 2 in the same dilution (λ490 of sample/λ490 of negative control). The analyses were carried out in duplicate.

### Cells and virus used in neutralization assays

African green monkey kidney (Vero, subtype E6) cells were cultured at 37°C in high-glucose DMEM with 10% fetal bovine serum (HyClone/Cytiva, USA), 100 U/mL penicillin and 100 μg/mL streptomycin (ThermoFisher, USA) in a humidified atmosphere with 5% CO_2_.

SARS-CoV-2 was prepared in Vero E6 cells. Originally, the isolate was obtained from a nasopharyngeal swab from a confirmed case in Rio de Janeiro, Brazil (IRB approval 30650420.4.1001.0008). All procedures related to virus culture were handled in a biosafety level 3 (BSL3) multiuser facility according to WHO guidelines. Virus titers were determined as plaque-forming units (PFU)/mL. The virus strain was sequenced to confirm the identity, and the complete genome is available in GenBank (SARS-CoV-2/human/BRA/RJ01/2020, #MT710714). The virus stocks were kept at −80°C.

### Microneutralization assay

To access the neutralization titer, the samples were incubated with 100 PFU of SARS-CoV-2 for 1 h at 37°C. Then, the samples were transferred to 96-well plates with monolayers of Vero cells (2 × 10^4^ cells/well) with serial dilutions of sample for 1 h at 37°C. Cells were washed, and fresh medium with 2% FBS and 2.4% CMC was added. On day 3 post-infection, the cytopathic effect was scored in at least 2 replicates per dilution by independent readers. The readers were blind with respect to the sample ID.

Statistical analyses were performed with GraphPad Prism 7®. Neutralization assay data are shown as the mean ± SD; p<0.05 vs. plasma collected on day 0 according to Student’s t-test for paired samples.

### Neutralizing titers in plasma of mice injected with equine F(ab’)_2_

In order to investigate if neutralizing activity against SARS-CoV-2 can be observed in animals treated with the fragments, equine F(ab’)_2_ (pilot lot) was injected by the intraperitoneal route and neutralizing titers were measured in the plasma of mice. 11 Balb/C mice were divided into 4 groups, which received different doses of the antibody fragments: 50 μg (n = 3); 25 μg (n = 3); 10μg (n = 3) and negative control (saline) (n = 2). The total volume injected was 200 μL per mouse. Blood was collected from the retro-orbital plexus 72 hours after inoculation, and then the animals were euthanized by saturation with isoflurane. The neutralization titer in mice plasma was determined by the plate reduction neutralization test (PRNT). Statistical significance was calculated using Graphpad Prism® 7 software, by one-way ANOVA and Tukey post-hoc test to the confidence levels indicated in Figure S2.

### Sample collection from human subjects

Samples of human convalescent plasma collected at the State Hematology Institute Hemorio followed a protocol approved by the local ethics committee (CEP Hemorio; approval #4008095), as described previously^15^.

## ACKNOWLEDGMENTS

The authors gratefully acknowledge B. S. Graham and K. S. Corbett (Vaccine Research Center, NIAID/NIH) for sharing the plasmid (BEI Resources # NR-52563) that enabled the generation of the cell line producing SARS-CoV-2 spike protein. We also thank Dr. A. M. Vale from UFRJ and Dr. S. Garcia and Dr. L. Amorim from Hemorio for human convalescent plasma samples. Financial support from Senai, CTG, Embrapii, Sebrae and from the Brazilian research funding agencies Fundação Carlos Chagas Filho de Amparo à Pesquisa do Rio de Janeiro (FAPERJ), Conselho Nacional de Desenvolvimento Científico e Tecnológico (CNPq), Coordenação de Aperfeiçoamento de Ensino Superior (CAPES) e Financiadora de Estudos e Projetos (FINEP).

## SUPPLEMENTARY MATERIAL

**Table S1.** Microneutralization data for equine sera collected during the first immunization of the first animal group, for F(ab’)_2_ concentrates and for human convalescent plasma.

| Sample ID |  | PRNT<sub>50</sub> | Mean±SD | PRNT<sub>90</sub> | Mean±SD |
| --- | --- | --- | --- | --- | --- |
| Equine preimmune serum | 832 | <16 | 16±0 | <16 | 16±0 |
|  | 833 | <16 |  | <16 |  |
|  | 835 | <16 |  | <16 |  |
|  | 841 | <16 |  | <16 |  |
|  | 842 | <16 |  | <16 |  |
| Equine serum on day 28 | 832 | 8,000 | 3,660±2,947 | 4,096 | 1,491±1,655 |
|  | 833 | 4,096 |  | 2,048 |  |
|  | 835 | 64 |  | 32 |  |
|  | 841 | 2,048 |  | 256 |  |
|  | 842 | 4,096 |  | 1,024 |  |
| Equine serum on day 35 | 832 | 16,000 | 19,302±13,203 | 8,000 | 9,651±6,601 |
|  | 833 | 32,000 |  | 16,000 |  |
|  | 835 | 512 |  | 256 |  |
|  | 841 | 32,000 |  | 16,000 |  |
|  | 842 | 16,000 |  | 8,000 |  |
| Equine serum on day 42 | 832 | 32,000 | 22,809±13,516 | 16,000 | 11,251±7,055 |
|  | 833 | 16,000 |  | 8,000 |  |
|  | 835 | 2,048 |  | 256 |  |
|  | 841 | 32,000 |  | 16,000 |  |
|  | 842 | 32,000 |  | 16,000 |  |
| Equine serum on day 49 | 832 | 32,000 | 23,219±12,738 | 32,000 | 14,604±11,560 |
|  | 833 | 16,000 |  | 8,000 |  |
|  | 835 | 4,096 |  | 1,024 |  |
|  | 841 | 32,000 |  | 16,000 |  |
|  | 842 | 32,000 |  | 16,000 |  |
| Equine F(ab')<sub>2</sub> concentrate | Pilot lot | 32,000 |  | 16,000 |  |
| Plasma pool used in GMP production (1<sup>st</sup> immunization of Groups 1 and 2) | Pool of 2 bleedings (160 L) | 16,384 |  | 4,096 |  |
| GMP equine F(ab')<sub>2</sub> concentrate | Industrial lot (16 L API) | 65,556 |  | 16,384 |  |
| Human convalescent plasma | 1 | 512 | 298±195 | 128 | 106±36 |
|  | 2 | 128 |  | 64 |  |
|  | 3 | 256 |  | 128 |  |

**Figure S1.**
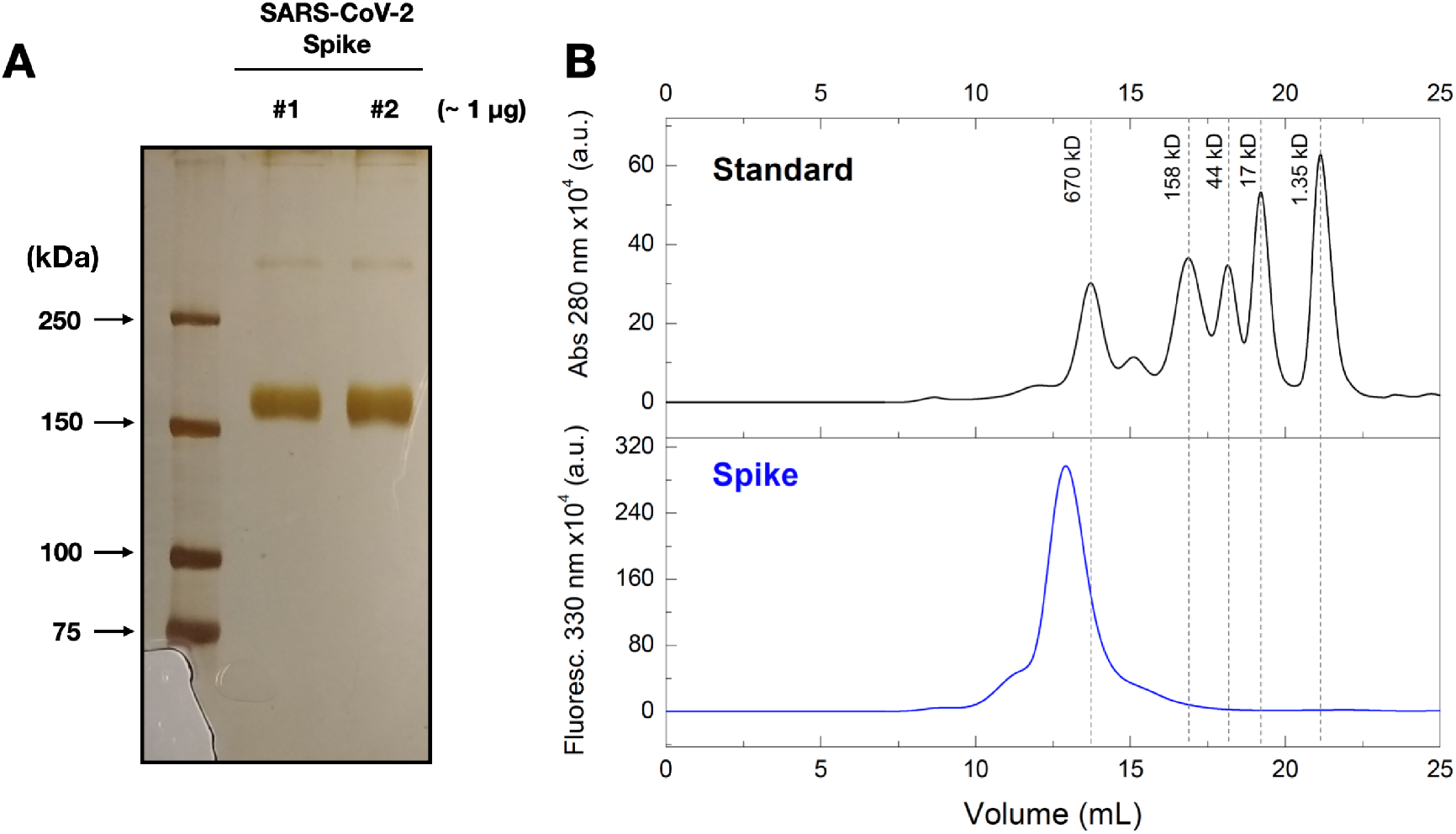
Quality checks of SARS-CoV-2 spike protein. **(A)** Image of a silver-stained 7% SDS-PAGE gel showing two representative batches of SARS-CoV-2 spike preparation (two batches for purposes of comparison). **(B)** Line plots showing size-exclusion chromatograms using a Superose 6 10/300 column. The fluorescence intensity at 330 nm of the spike protein was recorded as a function of the retention volume. The absorbance at 280 nm of a molecular mass standard (black line) is shown for comparison.

**Figure S2:**
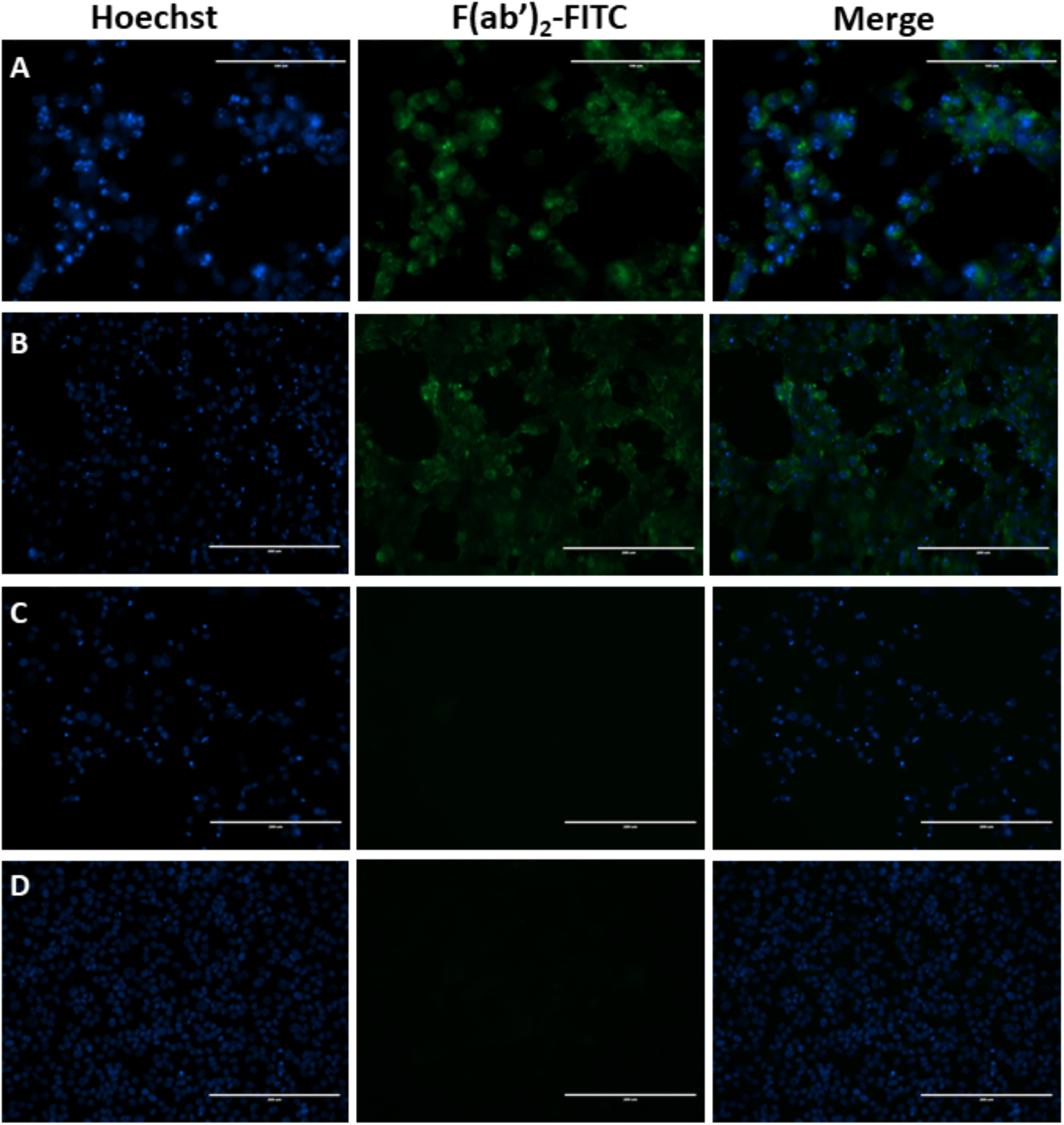
Equine F(ab’)_2_ binds specifically to SARS-CoV-2 infected cells. Vero E6 cells were infected with SARS-CoV-2 (MOI = 0.05) for 48 h and immunolabeled with F(ab’)_2_-FITC conjugate. **(A)** Cells infected with SARS-CoV-2, labeled with F(ab’)_2_ -FITC, and observed with 40x objective. **(B)** Cells infected with SARS-Cov-2 and labeled with F(ab’)_2_-FITC, and observed with 20x objective. **(C)** Control infected cells incubated in the absence of F(ab’)_2_-FITC. **(D)** Control infected cells incubated in the presence of non-labeled F(ab’)_2_. First column shows images in the blue channel corresponding to Hoechst labeling of cell nuclei, second column shows images obtained in the green channel for the detection of F(ab’)_2_-FITC fluorescence, and third column is a merge of blue and green channels. Bars on (A) 100 μm, on all other images 200 μm.

